# Shell accumulation and taphonomic attributes of the stout razor clam *Tagelus plebeius* as indicator of environmental features in a South American coastal lagoon of international relevance

**DOI:** 10.1101/2025.07.11.664465

**Authors:** Daniela M. Bernat, Guido L. Bacino, Paula A. Cristini, Mariana S. Addino

## Abstract

The preservation of remains is conditioned by biotic and abiotic characteristics that are determined by the environmental conditions; thus, taphonomic analysis may indicate environmental features. Estuaries are priority areas due to their socio-ecological importance; however, they are at risk due to climate change and anthropogenic impacts. Bivalves are abundant in estuarine soft-bottoms and their remains may be valuable environmental indicators, since the biotic (e.g., life habits, mineral composition) and abiotic characteristics (e.g., tides or topography) lead to differences in the accumulation and preservation of their shells. Therefore, the objective of this work was to evaluate the accumulation and taphonomic aspects of modern shells of the stout razor clam *Tagelus plebeius* in relation to the topographic characteristics of intertidal areas in Mar Chiquita coastal lagoon (Buenos Aires, Argentina). The results would indicate remains transport to the intertidal areas with a low-energy dynamic. The topographic characteristics of the beaches combined with the pattern of shell accumulation, suggested that the remains transport is not due directly to topography but to a combination of factors. In fact, those factors related to beach localization with respect to the main water flow direction and hydrodynamics would be more relevant. Furthermore, the findings showed that the taphonomic attributes of deposited modern shells may indicate environmental characteristics, such as energy of transport, submersion time or predation pressure. This study showed the contribution that actualistic taphonomy research can make to understanding environmental characteristics and to diagnostics development for the management of these vulnerable and socio-economic important areas.

**Statements and Declarations:** There is no competing interests related to this publication.

## 1. Introduction

The preservation of remains is conditioned by ecological and evolutionary processes that occur continuously and are usually determined by the particular biotic and abiotic characteristics of different environments; thus, the sedimentary record is rich in biological data and ecological information. Both the recent and distant records can be used to understand the responses of environments and species to long-term environmental changes. The shells of mollusks, particularly bivalves, which in modern estuaries tend to represent the largest proportion of the biomass in soft-bottom sediments (e.g., Seitz *et al*., 2006), are biological archives that provide temporal records of environmental conditions (e.g., Addino *et al*., 2024). The interpretation of mollusk shells to reconstruct the depositional environment depends on different factors. On one side, intrinsic biological characteristics, such as life habits, organic content and shell^′^s mineralogy, influence their susceptibility to damage and preservation (Lockwood and Work, 2006). On the other side, shell breakage is well-known in high-energy environments, through dissolution, abrasion and transport after death (Zuschin and Stanton, 2001), but it is also common in low-energy habitats where biological interactions, such as predation and bioturbation, are an important source of fragmentation.

Likewise, the morphology and orientation of coastline determines other aspects of depositional processes. Variations such as topographic relief and tidal range result in significant variations in the distribution and abundance of shell banks within shallow marine sediments (e.g., Meldahl, 1993). Both the duration of submersion and exposure to hydrodynamic energy, which are influenced by topography, create variations in shell accumulation (e.g., Schneider-Storz *et al*., 2008). Additionally, differences in current dynamics, tides, waves, and winds lead to differences in the accumulation and composition of the deposited material in dune barriers (e.g., Tsolakos *et al*., 2021). Therefore, numerous and combined environmental characteristics affect taphonomic processes, so that the taphonomic features found in the remains have the potential to reveal environmental processes on different temporal scales.

Estuaries are considered priority areas due to their ecological importance and their economic contribution to coastal areas (see Barbier *et al*., 2011). They are highly productive areas and provide a variety of ecosystem services, such as acting as buffer zones against storm and flood events (Leonardi *et al*., 2018) or providing a special site for recreation and tourism (Barbier *et al*., 2011). Furthermore, they are extremely unstable ecosystems due to their limited connection to the sea, their shallow depths that are highly sensitive to changing climatic conditions, the role of human interventions, and finally, their characterization by low biodiversity and high productivity (Elliot *et al*., 2014). Globally, estuaries have been documented to be at risk due to the effects of climate change and anthropogenic impacts. In particular, the accelerated sea level rise would imply a series of effects on these ecosystems, whose consequences are currently difficult to predict.

In estuaries, the particular role of tides in altering the water level causes a daily (or bi-daily) change in flow direction and directly affects sediment dynamics. For example, tides shape intertidal shoals and tidal channels, which can modify the effects of waves on beach formation (Vila-Concejo *et al*., 2020). These low energy beaches, controlled by locally generated wind-waves (fetch-limited) are characterized by a narrow or absent backshore, steep foreshore and a subaqueous large low-tide terrace (Jackson, 1995). Its morphology is mostly affected by short term variations in wave energy caused by storms, for this reason the observed morphology may be a relic feature related to the last high-energy event, since post-storm low intensity average conditions (high frequency) may not be sufficient to return beaches to their original conditions (Jackson et al., 2002).

The stout razor clam *Tagelus plebeius* is an euryhaline, filter-feeder infaunal bivalve that inhabits estuarine tidal flats with cohesive sand and silt sediments. It is present along the American Atlantic coast from Cape Cod, Massachusetts (42° N, USA) to northern argentinean Patagonia (41° S). In the northern coast of Argentina, it inhabits at least four estuaries with very different environmental settings, one of these is Mar Chiquita coastal lagoon (37°33^’^ S - 57°30^’^ W). This is the only coastal lagoon associated with a currently active litoral barrier in Argentina, making it a significant area due to its complexity and socio-ecological importance. It holds protected status at multiple levels of government (Local, Provincial, National), as well as internationally (UNESCO^’^s MAB Program). *T. plebeius* is the only native bivalve species inhabiting the intertidal flats of these estuaries, living buried in permanent burrows (Addino *et al*., 2016; Addino *et al*., 2019). These clams are the main prey item of the American oystercatcher *(Haematopus palliatus*; Bachmann and Martínez, 1999) and it is an important component of the infauna that plays an important role in various ecological processes (e.g., Gutierrez *et al*., 2003; Addino *et al*., 2015; Addino *et al*., 2019; Alvarez *et al*., 2015) and, consequently, in the habitat functioning.

Moreover, within the shell ridges deposited along the South Atlantic coast as a result of Holocene sea level fluctuations, well-defined horizons containing articulated shells of *T. plebeius* in life position are common (e.g., Aliotta and Farinati, 1990). They may be explained by oystercatcher predation (Iribarne *et al*., 1998) and gradual accumulation under stable conditions (Golfieri *et al*., 1998). These are of particular interest for paleoenvironmental studies of the Southwestern Atlantic, as they are widely distributed from southern Uruguay to northern Argentine Patagonia (Aguirre and Fucks, 2004; Martínez and Del Río, 2005) and they are reliable indicators of past estuarine environments that developed during the last transgressive-regressive cycle (Addino *et al*., 2024). Additionally, reworked fossil deposits along with remains from modern populations of these clams have been found, and their characteristics hold implications for environmental reconstruction (De Francesco and Hassan, 2008). As it was mentioned above, the deposition of these remains may be related to environmental drivers such as topography or hydrodynamics. In this context, the objective of this work was to evaluate the accumulation and taphonomic aspects of modern *Tagelus plebeius* shells in relation to the topographic characteristics of intertidal areas in Mar Chiquita coastal lagoon (Buenos Aires, Argentina), in order to determine its usefulness and future projection as an analytical tool for the integrated study of coastal systems.

## 2. Materials and methods

### 2.1. Study area

This study was conducted in Mar Chiquita coastal lagoon (Fig. 1), Buenos Aires province, Argentina. This lagoon covers an area of approximately 46 km^2^, with a north-south elongated shape, permanently connected to the sea (Isla, 1997). The entrance channel is approximately 6 km long, 200 m width and its depth is between 1.5 and 2 m (Fasano *et al*., 1982; Reta *et al*., 2001). It is characterized by the influence of a semidiurnal tidal regime with a microtidal amplitude (0.8 m; Isla and Gaido, 2001), where winds and rainfall significantly alter the water level variations (Isla and Garrido, 2001). Its geomorphology was originated during the Holocene in response to the interaction between sea-level fluctuations and various coastal dynamic factors, which transformed this semi-enclosed coastal basin into one of the main estuarine depocenters, where low-energy conditions prevailed due to its isolation from the open sea (Violante and Cavallotto, 2004). The specific study area was set in the inlet channel of the coastal lagoon. Particularly, in the intertidal area of four artificial beaches, hereafter referred to as “beaches”, where preliminary observations indicated some geomorphological differences among them. They are delimited by artificial breakwaters of 20 m length, spaced approximately 140 m and located 700 m from the inlet mouth to the sea (Fig. 1).

**Fig.1:**
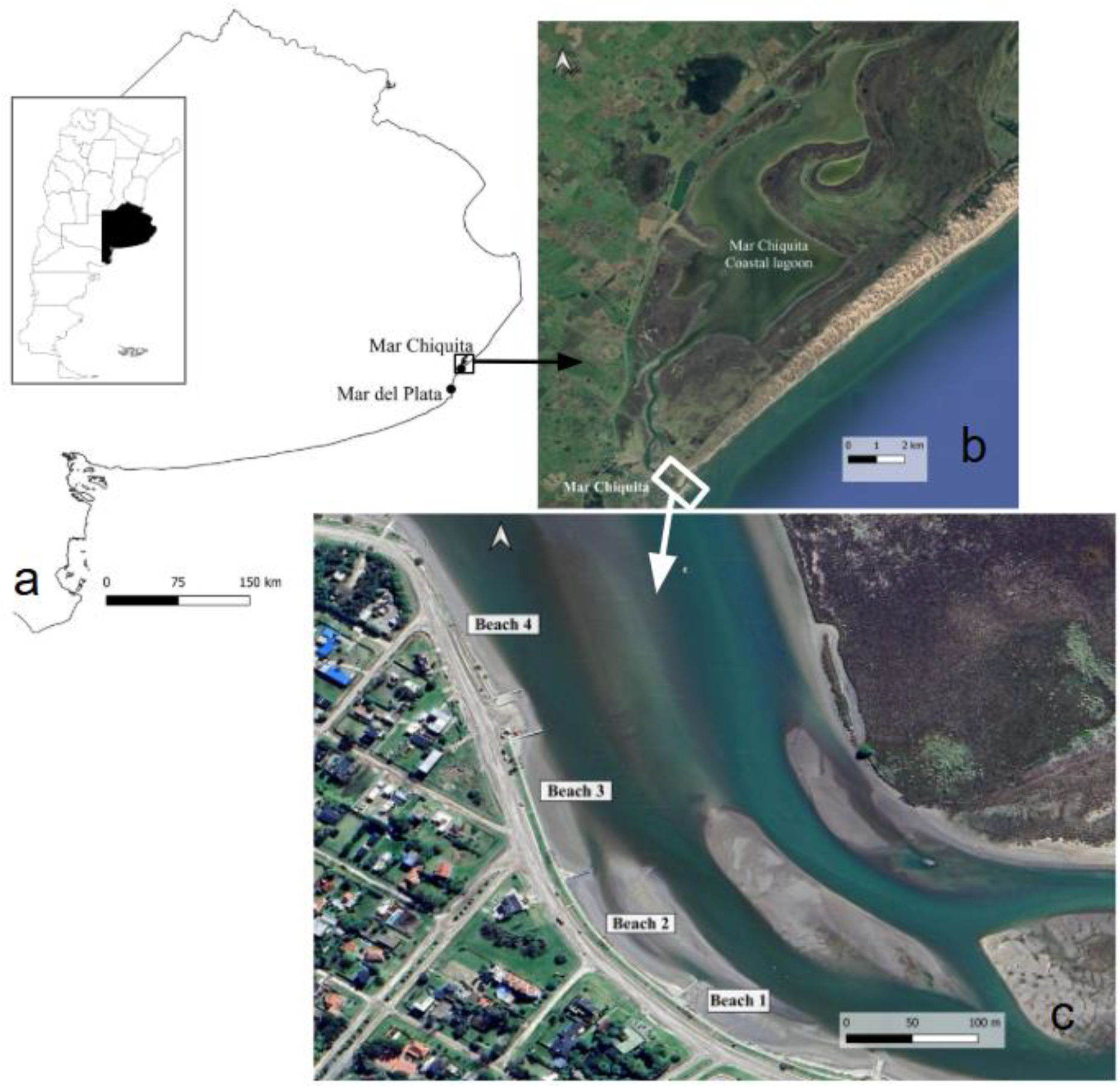
a) Geographic location of the study area into Buenos Aires province (Argentine), b) Mar Chiquita coastal lagoon, and c) Location of the sampling sites (beaches).

### 2.2. Shell accumulation in the beaches

A transect was drawn from the beginning to the end of each beach (i.e., from one breakwater to the next one), parallel to the shoreline on the foreshore (i.e., where all high tides reach and it is uncovered in each low tide). Along each transect, a 1m diameter sampling unit was established every 10 linear meters. In each sampling unit the presence of articulated shells of *T. plebeius* was registered and, in order to perform a more powerful analysis, this procedure was repeated in four dates (between August and November 2023). The percentage of sampling units with articulated shells was compared between beaches with a *Chi-square* test for each date. Given that there was the same pattern of shell presence in the different beaches for each of the four dates (see Results), the temporal variation could be discarded and therefore, samplings by date were pooled into each beach for further analyses. Then, the proportion of sampling units with “presence” of shells was compared between beaches.

Additionally, during the first sampling date, all shells found within an area of one meter to each side of the transect were collected for further taphonomic analysis (see 2.3).

### 2.3. Taphonomic analysis

The collected shells of *T. plebeius* were taken to the laboratory, where they were washed, measured in anterior-posterior length using a calliper (hereafter Length, precision 0.05 cm), when at least one valve was complete, and subjected to taphonomic observations. Length was compared between beaches using the Kruskal-Wallis test followed by the Wilcox test (Zar, 1999), since the data did not show a normal distribution (see Results). The proportions of shells with and without: 1) surface alteration (loss of the periostracum on more than 20% of the shell, hereafter ^“^Periostracum^”^), 2) ^“^Fragmentation^”^ (i.e. shells with more than 20% of their structure broken), and 3) presence of predation marks by oystercatchers (breakage in the posterior part of the shell (Lomovasky *et al*., 2005), hereafter ^“^Predation^”^) were established (Fig. 2) and compared between beaches by means of *Chi-square* tests.

**Fig.2:**
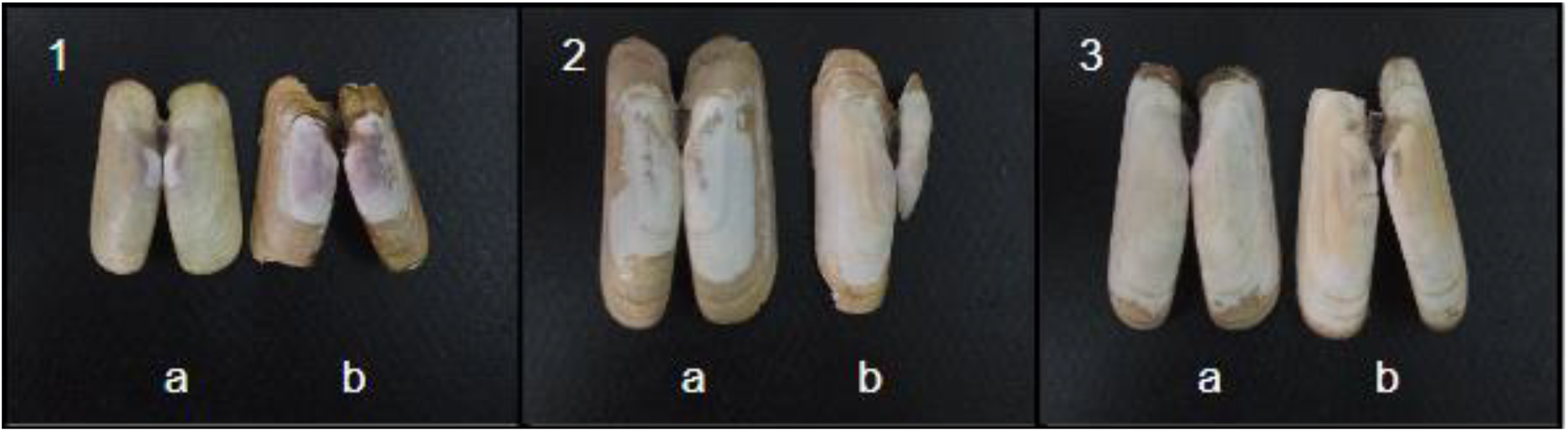
Shells with the different taphonomic attributes considered in this study as dichotomous variables: 1.a) Not Altered Shell (NA), 1.b) Altered Shell (A), 2.a) Not Fragmented Shell (NF), 2.b) Fragmented Shell (F), 3.a) Shell Without Predation Mark by oystercatchers (SM), and 3.b) Shell with Predation Mark (M, breakage in the posterior part)

### 2.4. Topographic data

The Structure from Motion Multi-View Stereo technique (Westoby *et al*., 2012) was used to reconstruct topography from images. These images were captured using a DJI Phantom Pro drone equipped with a 1-inch sensor camera. The flights were programmed autonomously with the DroneDeploy application (https://www.dronedeploy.com/), at an altitude of 40 m with a 70% front and side overlap between images. In the flight, conducted in July 2022, control points were measured to georeference and adjust the model, using differential GNSS with two Emlid Reach Rs2 antennas. A total of 25 points were measured, 5 of which were fixed points. With the CloudCompare software (version 2.7), the point cloud was segmented by beach and filtered to remove elements outside of the beach, using the Cloth Simulation Filter (CSF) tool. Further analysis where the shell sampling transect was conducted, the intertidal zone (foreshore), was isolated in order to determine the average slope, the average elevation and the beach orientation.

## 3. Results

### 3.1. Shell accumulation in the beaches

The beaches displayed a similar pattern across different sampling dates, with beaches 3 and 4 reaching in general a higher or equal proportion of sampling units with presence and absence of shells (Fig. 3); thus the samplings by dates were pooled for each beach.

**Fig.3:**
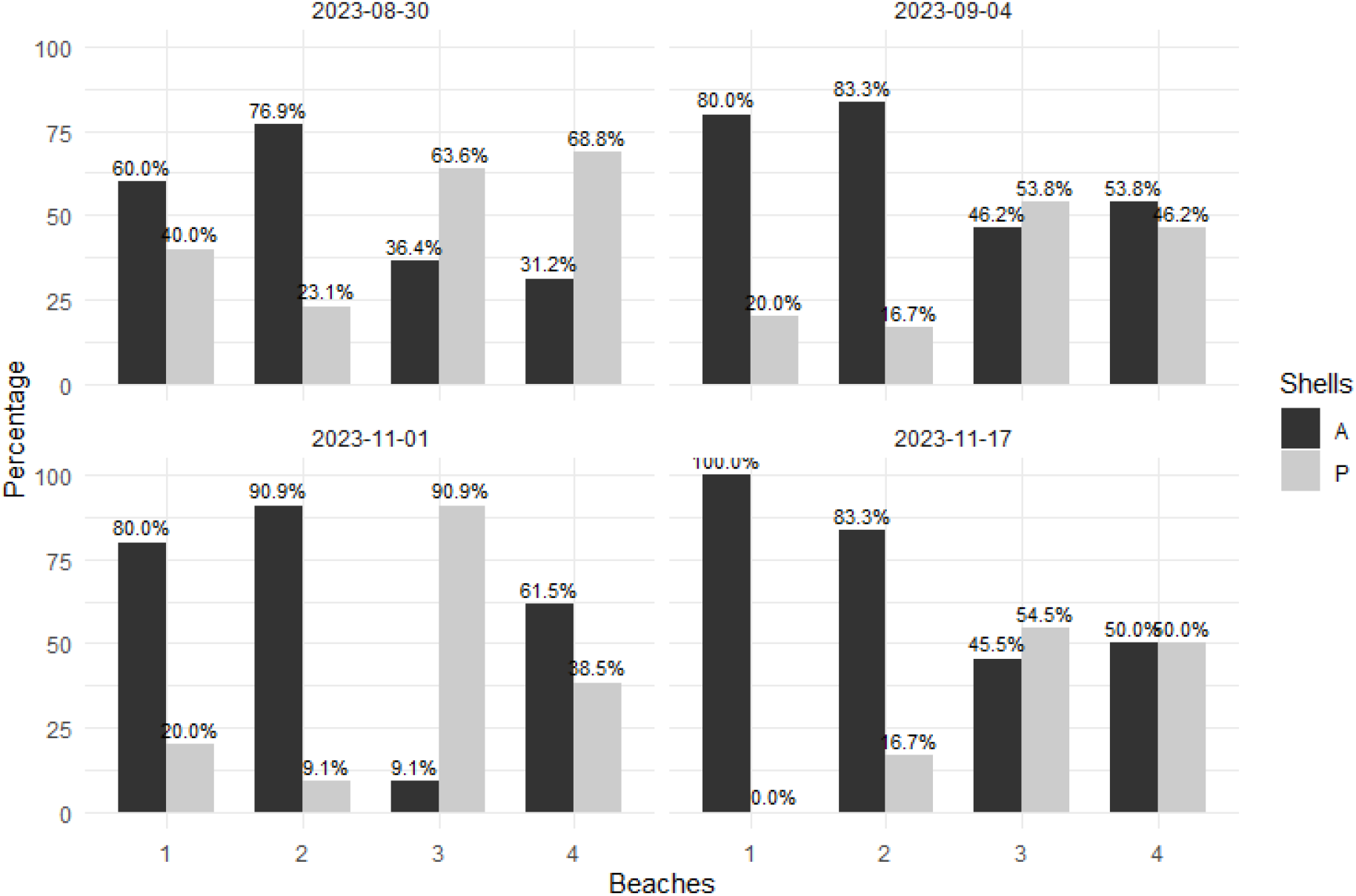
Percentage of sampling units with Presence (P) in grey and Absence (A) in black of shells, recorded at each beach on different sampling dates

The *Chi-square* test showed significant differences for the shell^’^s presence proportion between the beaches (Χ^2^= 30.263; p < 0.05), with the beaches 3 and 4 showing the highest values of presence (Fig. 4).

**Fig.4:**
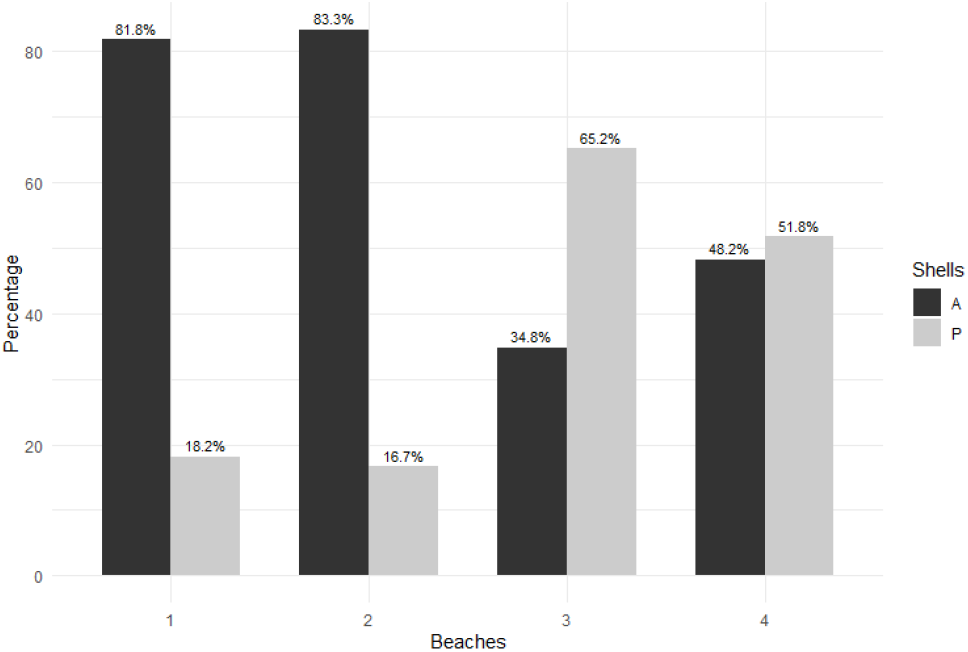
Percentage of sampling units with Presence (P, in grey) and Absence (A, in black) of shells on each beach. 100% represents the total sampling units on each beach

### 3.2. Taphonomic analysis

The number of shells collected for taphonomic analysis varied across beaches, with only 3 shells collected from beach 1, 46 from beach 2, 201 from beach 3, and 213 from beach 4. Due to the lack of representativeness, the shells from beach 1 were excluded from the analysis. Length distribution was not normal (Shapiro-Wilk: B2: W= 0.890; B3: W = 0.885; B4: W = 0.817; p < 0,05 for all beaches). The average Length of the collected shells showed significant differences between beaches (Kruskal Wallis test: Χ^2^ = 31.396; p < 0.05). Results from the Wilcox test indicated that shells from beach 2 are smaller than those collected from beaches 3 and 4. Meanwhile, shells from beaches 3 and 4 do not show significant differences between them (Fig. 5).

**Fig.5:**
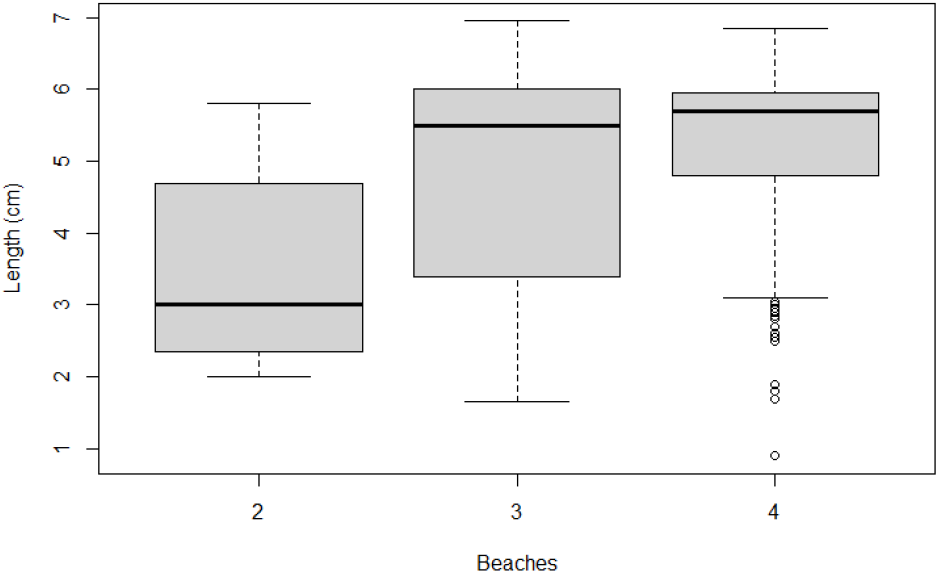
Anterior-posterior length of the shells collected at each beach. The lower and upper boundaries of the boxes represent the 25% and 75% percentiles, respectively, the horizontal lines within the boxes indicate the median value, while the limits of the vertical lines show the maximum and minimum values, and the points represent outliers

*Chi-square* test showed significant differences for the variables Periostracum (Χ^2^ = 68.014; p < 0.05) and Fragmentation (Χ^2^ = 27.133; p < 0.05) among beaches. Beaches 3 and 4 showed a higher proportion of shells with altered periostracum (Fig. 6) and a lower proportion of fragmented shells (Fig. 7) than beach 2. On the other hand, the proportion of shells with predation mark did not show differences between beaches (Χ^2^ = 0.012; p > 0.05, Fig. 8).

**Fig.6:**
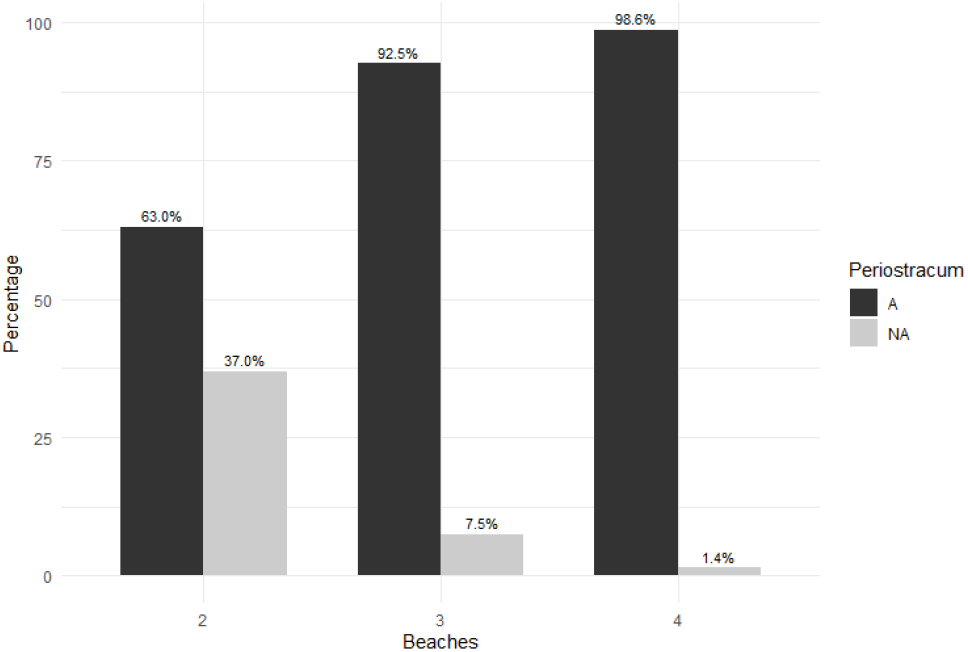
Percentage of shells with altered periostracum (A, in black) and not altered periostracum (NA, in grey) in the different beaches. 100% represents the total number of shells on each beach.

**Fig.7:**
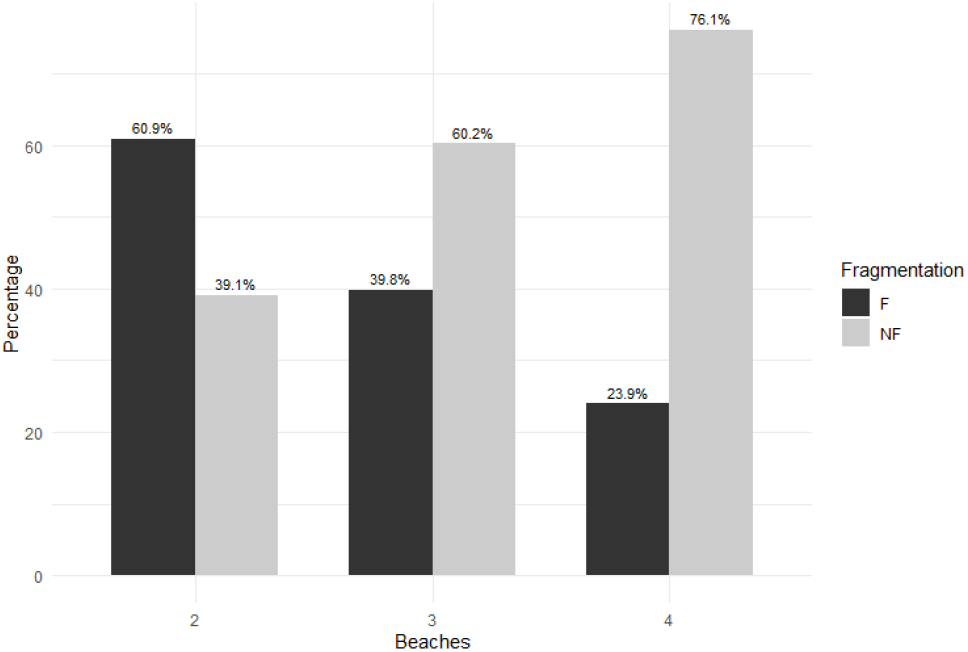
Percentage of shells fragmented (F, in black) and not fragmented (NF, in grey) in the different beaches. 100% represents the total number of shells on each beach.

**Fig.8:**
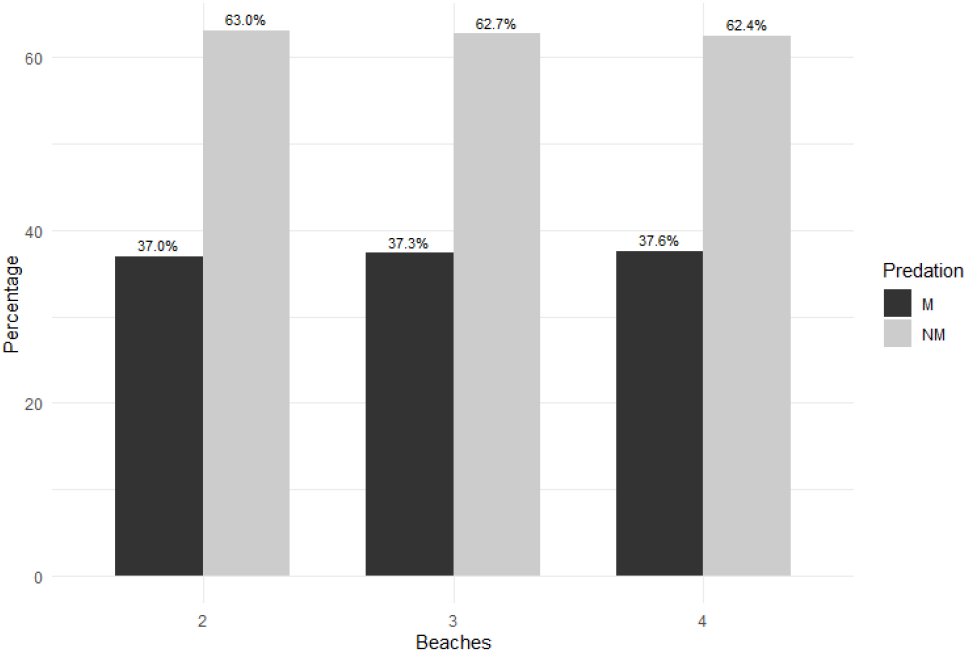
Percentage of shells with predation mark (M, in black) and without predation mark (NM, in grey) in the different beaches. 100% represents the total number of shells on each beach

### 3.3. Topographic data

The 3D point cloud models resulted with a Root Mean Square Error (RMSE) of the control points ranging between 0.21 m and 0.36 m, and the reprojection error of the models had values between 0.11 m and 0.15 m, indicating that the geometry of the beach surface could be represented with sufficient precision (James *et al*., 2017).

The four evaluated beaches exhibited two different patterns in terms of their shapes and widths. Beach 2 and 4 are characterized by their narrower profiles, with an average width of 15 m and an average slope of 3.5°, whereas beaches 1 and 3 displayed wider profiles, averaging 30 m in width and featuring gentler slopes averaging 1° (Fig. 9). It is worth noting that beach 1 shows greater variability in the elevation and slope due to the discharge of rainwater pipe, which has altered its morphology. Respecting the specific shell sampling sector of each intertidal zone (white polygons in fig 9), beach 3 resulted with the lowest average slope estimated (0.64°) meanwhile beaches 1 and 2 resulted with similar slopes (1.72 - 1.80°) and beach 4 resulted with the highest slope average, 2.18°. On the other side, beach 4 resulted with the highest height above mean sea level (0.22 m) followed by beach 3 (−0.2 m). Beaches 1 and 2 showed the lowest elevation (−0.34 m). For the orientation relative to the North, beaches 1 and 3 have the greatest angle; i.e., they are more oriented toward the East (Fig. 9, Table 1).

**Table 1:** Average slope values, elevation, and orientation in each beach intertidal transect. Slope values are expressed in degrees (°) and as β (where β = tan(angle)). Elevation is referenced to the IGN datum and the orientation refers to the perpendicular to the beach and is relative to North (0°).

|  | Beach 1 | Beach 2 | Beach 3 | Beach 4 |
| --- | --- | --- | --- | --- |
| Slope ($^{\circ}/\beta$) | 1.80 / 0.031 | 1.72 / 0.03 | 0.64 / 0.011 | 2.18 / 0.038 |
| Elevation (m) | -0.34 | -0.34 | -0.2 | 0.22 |
| Orientation ($^{\circ}$) | 81 | 59 | 85 | 63 |

**Fig.9:**
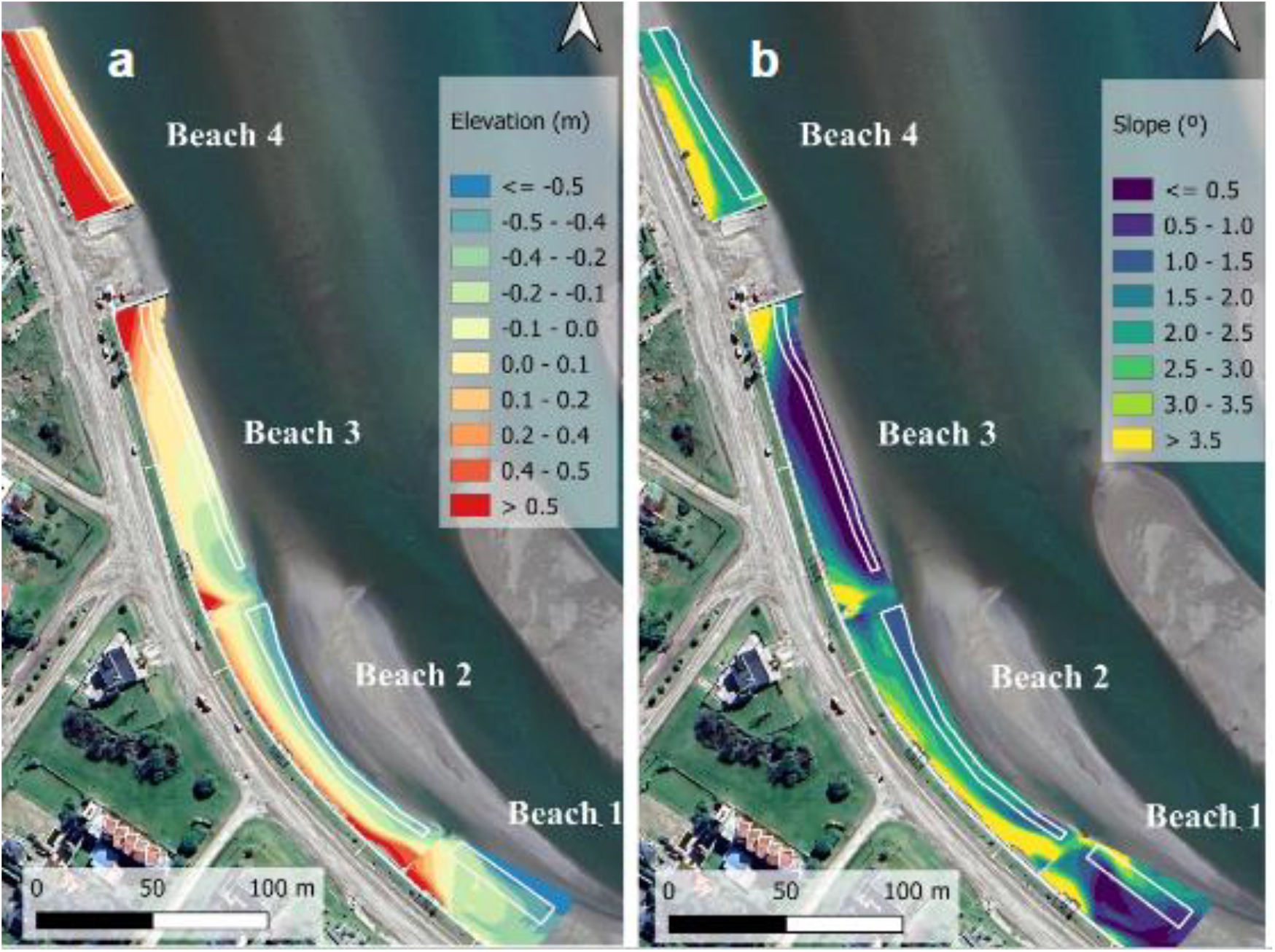
a) Digital elevation models (in meters) and b) the slope (in degrees), of the analysed beaches. The intertidal transects, where average values were computed, are indicated in the white polygons.

In addition, the digital elevation model allowed for the visualization of a sandbank in front of beaches 1 and 2, as well as a tidal channel between this sandbank and those beaches. Also greater accumulation of sediment can be observed on the northwest side of the breakwaters (Fig. 9).

## 4. Discussion

The results of the present study indicate that the pattern of shell accumulation in the beaches did not change over time, showing that beaches 3 and 4 presented a higher probability of finding *Tagelus plebeius* shells on the sediment. Taphonomic analyses showed that shell length distribution was not normal. Given that in the living population of *T. plebeius* shell length is normally distributed (Addino *et al*., 2019), this finding could suggest an effect of shell transport. Furthermore, beach 2 had smaller shells compared to the other beaches. This can also result from a differential transport of shells to this beach, determined by changes in hydrodynamic energy due to the presence of a sandbank in front of beaches 2 and 1 (Fig. 9).

Baches 3 and 4 had a higher proportion of shells with altered periostracum and a lower proportion of fragmented shells. Meanwhile, shells in beach 2 also presented a high proportion of periostracum alteration but with a higher proportion of fragmentation. The presence of the periostracum in shells is believed to protect against acid dissolution (Saleuddin and Petit, 1983). This protective layer tends to degrade after prolonged submersion under a film of water with suspended fine material, which acts as an abrasive on the shell surface (Pereira *et al*., 2021). Thus, taphonomy analysis suggests that the shells on beaches 3 and 4 experienced prolonged submersion and low transport, i.e., low fragmentation. In contrast, on beach 2, the effect of transport energy may be greater generating shell fragmentation. This is consistent because this beach has the lowest mean intertidal elevation of -0.34 m (Table 1) and also reinforces prior suggestions.

The record of shells with bird predation mark was around 37 percent, i.e. lower than without mark in all the beaches, which is consistent with that reported by Cadeé (1995) for the Wadden Sea, where 30-35% of carbonate remains were generated by bird predation. Likewise, the low percentage of shells with predation marks would indicate that the production and accumulation of remains is due to a combination of factors, including transport, as it was mentioned before. Overall, these results suggest a low-energy dynamic, a typical characteristic of estuarine beaches (Davis and Fitzgerald, 2009), but showing that the different taphonomic attributes (length, surface alteration, fragmentation and predation mark) may indicate site-specific environmental characteristics.

The higher shell accumulation in beaches 3 and 4 did not show an evident correlation with topographic characteristics. Both beaches had a similar probability of finding *Tagelus plebeius* shells but exhibited the lowest (0.64°) and the highest (2.18°) mean intertidal beach slope, respectively. Additionally, the mean elevation on beach 4 is 0.4 m higher than that of beach 3. Both beaches can be considered smooth in general (≤ 3°; Makaske and Augustinus, 1998). When beach face slope and profile shape are compared with the model of low energy beaches morphotype proposed by Travers (2007), beaches 1, 2 and 4 align well with the exponential beach morphotype (slope 1.8-2.1°), except for its beach widths (∼200m). This beach morphotype is characterized by its protection, with very low exposure to wind-waves caused by the short fetch distance. Together with the shell taphonomic characteristics, the topographic characteristics indicated low energy and prolonged submersion for the studied site, at least in beach 3 and 4. In the case of beach 2, where there is more shell fragmentation, it may be related to the changes in bathymetry and the hydrodynamics generated by the sandbank reducing swash effects, while enhancing channel current flow velocity, which also affect beach 1.

Orientation would be an important factor if the suggested effect of shell transport is occurring, since the main channel current comes from the N-NW sector where there are dense clam populations (Addino *et al*., 2019). In this context, it would be expected low remains arrival for beaches 1 and 3 since these are slightly more orientated to the East; however, the opposite pattern was found in beach 3, given that it had the highest probability of finding shells. Unlike oceanic beaches, in estuaries waves can reach the coastlines at high angles, inducing longshore currents and potential transport of bioclasts and fine sediments (Vila-Concejo *et al*., 2020). This, along with the current speed, which reaches the highest speeds in the study site with a maximum of 140 cm/s (Lanfredi *et al*., 1987), would indicate that the dominant factor in shell accumulation is the location of the beaches, not the orientation, relative to the direction of the current flow. Therefore, beach 3 and 4 would be the first to receive the transported shells. In fact, sediment accumulation on the northwest side of the beaches (Fig. 9) would demonstrate this deposition pattern. The similarity in slope and the difference in the probability of finding shell accumulation between beaches 1 and 2 and beach 4 would also support this, suggesting that the coastal hydrodynamic factor would have more influence on shell accumulation. The sandbank in front of beaches 1 and 2 and the tidal channel formed between the bank and these beaches could prevent shell deposition and generate more fragmentation during the transport.

The findings presented in this work showed that shell accumulation in the intertidal zones of Mar Chiquita coastal lagoon does not depend on a single factor but rather a combination of them; being the transport of remains and the environmental features that affect it the most relevant. The high shell accumulation on beach 4 supports the suggestion that the specific flow hydrodynamics of the lagoon access channel may be more relevant than the beach morphometric characteristics such as slope or elevation. At the same time, the results obtained allow to identify that, at least in high-accumulation sites like beaches 3 and 4, the taphonomic variables can indirectly indicate the environmental energy determined by topographic variables, such as slope or elevation, and hydrodynamic variables, such as the main channel current direction. Furthermore, the lower accumulation of remains on beaches 1 and 2 highlighted the need to evaluate the hydrodynamic factor itself and in relation to geomorphology, as it can be affected by surrounding geoforms such as the observed sandbank in front of those beaches.

The results of this study provide a novel contribution to the field of actualistic taphonomy in the region and its approach in estuaries, considering that in these environments knowledge is limited, despite the recent growth of the discipline in the region (see De Francesco *et al*., 2020). Moreover, this study encourages further research on beach morphometrics and nearshore hydrodynamics in relation to their effects on shell taphonomy across different beach types. It is known that the rapid rise in sea level would imply a series of negative effects on these ecosystems, such as flooding and erosion (Silvestri *et al*., 2018). The contribution that actualistic taphonomy can make to understanding environmental characteristics, as demonstrated here, suggests the usefulness of this discipline in generating diagnostics which are necessary for the management of these maritime-front environments under potential vulnerability.

## 5. Conclusion

The accumulation of shells of the stout razor clam *Tagelus plebeius* in the intertidal zones of Mar Chiquita coastal lagoon appears to be related to the transport of the remains, which is affected in some beaches by the topographic characteristics, while in others it seems to be related to coastal hydrodynamics. The taphonomic attributes of the deposited shells were indicative of environmental characteristics, such as the energy of the environment. In this regard, it was shown that actualistic taphonomy is a powerful tool for identifying environmental attributes that may be particularly relevant for management in these areas, which are vulnerable to the current changes in sea level.

## Acknowledgements

This study was partially founded by the Facultad de Ciencias Exactas y Naturales-Universidad Nacional de Mar del Plata (FCEYN-UNMDP) through the research projects EXA1114-22 and EXA1175/24, and a research training fellowship to Daniela M. Bernat from the Consejo de Investigaciones Científicas of the Buenos Aires province (CICPBA). These findings are part of the undergraduate thesis submitted by D.M. Bernat in partial fulfillment of the requirements for the Bachelor^’^s degree in Biological Sciences.

## Bibliography

Abrahão, J. R., Cardoso, R. S., Yokoyama, L. Q., & Amaral, A. C. Z. (2010). Population biology and secondary production of the stout razor clam Tagelus plebeius (Bivalvia, Solecurtidae) on a sandflat in southeastern Brazil. Zoologia (Curitiba), 27, 54–64.

Addino, M. S., Alvarez, M. F., Brey, T., Iribarne, O., & Lomovasky, B. J. (2019). Growth changes of the stout razor clam Tagelus plebeius (Lightfoot, 1786) under different salinities in SW Atlantic estuaries. Journal of Sea Research, 146, 14–23.

Addino, M. S., Farenga, M. O., Martinell, J., Raniolo, A., & De Francesco, C. G. (2024). Linear morphometry and shell growth of the bivalve Tagelus plebeius as an indicator for salinity in Holocene South American estuaries. Lethaia, 57, 1–10.

Addino, M., Montemayor, D. I., Escapa, M., Alvarez, F., Valiñas, M., Lomovasky, B. J., & Iribarne, O. (2015). Effect of Spartina alterniflora Loisel on growth of the stout razor clam Tagelus plebeius. Journal of Experimental Marine Biology and Ecology, 463, 135–142.

Aguirre, M. L., & Fucks, E. (2004). Moluscos y paleoambientes del cuaternario marino en el sur de Entre Ríos y litoral bonaerense. En F. Aceñolaza (Ed.), Temas de la Biodiversidad del Litoral Fluvial Argentino (Miscelánea 12, pp. 55–70). INSUGEO.

Aliotta, S., & Farinati, E. (1990). Stratigraphy of Holocene sand-shell ridges in the Bahía Blanca Estuary, Argentina. Marine Geology, 94, 353–360.

Alvarez, M. F., Addino, M., Iribarne, O., & Botto, F. (2015). Combined engineering effects of clams and crabs on infaunal assemblages and food availability in intertidal systems. Marine Ecology Progress Series, 540, 57–71.

Bachmann, S., & Darrieu, C. A. (2010). Biología reproductiva del ostrero pardo (Haematopus palliatus) en el sudeste de la provincia de Buenos Aires, Argentina. El Hornero, 25(2), 75–84.

Bachmann, S., & Martínez, M. M. (1999). Feeding tactics of the American oystercatcher (Haematopus palliatus) on Mar Chiquita coastal lagoon, Argentina. Ornitología Neotropical, 10, 81–84.

Bacino, G. L., Dragani, W. C., & Codignotto, J. O. (2019). Changes in wave climate and its impact on the coastal erosion in Samborombón Bay, Río de la Plata estuary, Argentina. Estuarine, Coastal and Shelf Science, 219, 71– 80.

Barbier, E. B., Hacker, S. D., Kennedy, C., Koch, E. W., Stier, A. C., & Silliman, B. R. (2011). The value of estuarine and coastal ecosystem services. Ecological Monographs, 81, 169–193.

Behrensmeyer, A. K., Kidwell, S. M., & Gastaldo, R. A. (2000). Taphonomy and paleobiology. Paleobiology, 26(S4), 103–117.

Cadée, G. C. (1995). Birds as producers of shell fragments in the Wadden Sea, in particular the role of the Herring gull. Geobios, 28, 77–85.

Clay, R. P., Lesterhuis, A. J., Schulte, S., Brown, S., Reynolds, D., & Simons, T. R. (2014). A global assessment of the conservation status of the American Oystercatcher Haematopus palliatus. International Wader Studies, 20, 62–82.

Davis Jr, R. A., & FitzGerald, D. M. (2009). Beaches and coasts (2nd ed.). Blackwell Publishing.

De Francesco, C. G., & Hassan, G. S. (2008). Dominance of reworked fossil shells in modern estuarine environments: Implications for paleoenvironmental reconstructions based on biological remains. Palaios, 23(1– 2), 14–23. 10.2110/palo.2006.p06-124r

De Francesco, C. G., Tietze, E., Cristini, P. A., & Hassan, G. S. (2020). Actualistic Taphonomy of Freshwater Mollusks from the Argentine Pampas: An Overview of Recent Research Progress. In S. Martínez, A. Rojas, & F. Cabrera (Eds.), Actualistic Taphonomy in South America (Vol. 48, pp. 173–194). Springer. 10.1007/978-3-030-41664-3_9

Elliot, M., Cutts, N. D., & Trono, A. (2014). A typology of marine and estuarine hazards and risks as vectors of change: A review for vulnerable coasts and their management. Ocean and Coastal Management, 93, 88–99.

Golfieri, G. A., Ferrero, L., & Zárate, M. (1998). Tafonomía y paleoecología de Tagelus plebeius (Lightfoot, 1786) (Mollusca, Bivalvia) en sedimentos holocenos de río Quequén Grande, provincia de Buenos Aires, Argentina. Ameghiniana, 35(3), 255–264.

Gutiérrez, J., & Iribarne, O. O. (1999). Role of Holocene beds of the scout razor clam Tagelus plebeius in structuring present benthic communities. Marine Ecology Progress Series, 185, 213–228.

Gutiérrez, J., Jones, C., Strayer, D., & Iribarne, O. (2003). Mollusks as ecosystem engineers: the role of shell production in aquatic habitats. Oikos, 101, 79–90.

Holland, A. F., & Dean, J. M. (1977). The biology of the stout razor clam Tagelus plebeius: I. Animal-sediment relationships, feeding mechanism, and community biology. Chesapeake Science, 18, 58–66.

Iribarne, O., Valero, J., Martínez, M. M., Lucifora, L., & Bachmann, S. (1998). Shorebird predation may explain the origin of Holocene beds of stout razor clams in life position. Marine Ecology Progress Series, 167, 301–306.

Isla, F. I., & Gaido, E. S. (2001). Evolución geológica de la laguna Mar Chiquita. In O. Iribarne (Ed.), Reserva de bísfera Mar Chiquita: Características físicas, biológicas y ecológicas (pp. 1–319). Editorial Martín.

Jackson, N. L. (1995). Wind and waves: influence of local and non-local waves on mesoscale beach behavior in estuarine environments. Annals of the Association of American Geographers, 85(1), 21–37.

Jackson, N. L., Nordstrom, K. F., Eliot, I., & Masselink, G. (2002). “Low energy” sandy beaches in marine and estuarine environments: a review. Geomorphology, 48, 147–162. 10.1016/S0169-555X(02)00179-4

James, M. R., Robson, S., & Smith, M. W. (2017). 3-D uncertainty-based topographic change detection with structure-from-motion photogrammetry: precision maps for ground control and directly georeferenced surveys. Earth Surface Processes and Landforms, 42(12), 1769–1788.

Kowalewski, M., & Labarbera, M. (2004). Actualistic Taphonomy: Death, decay, and disintegration in contemporary settings. Palaios, 19(5), 423–427.

Lanfredi, N. W., Balestrini, C. F., Mazio, C. A., Schmidt, S. A. (1987). Tidal Sandbanks in Mar Chiquita Coastal Lagoon, Argentina. Journal of Coastal Research, 3(4), 515–520.

Leonardi, N., Carnacina, I., Donatelli, C., Ganju, N. K., Plater, A. J., Schuerch, M., Temmerman S. (2018). Dynamic interactions between coastal storms and salt marshes: A review. Geomorphology, 301, 92–107.

Lockwood, R., Work, L. A. (2006). Quantifying taphonomic bias in molluscan death assemblages from the upper Chesapeake Bay: patterns of shell damage. Palaios, 21(5), 442–450.

Lomovasky, B. J., Gutiérrez, J. L., Iribarne, O. O. (2005). Identifying repaired shell damage and abnormal calcification in the stout razor clam Tagelus plebeius as a tool to investigate its ecological interactions. Journal of Sea Research, 54(2), 163–175.

Marcovecchio, J., Freije, H., De Marco, S., Gavio, A., Ferrer, L., Andrade, S., Beltrame, O., Asteasuain, R. (2006). Seasonality of hydrographic variables in a coastal lagoon: Mar Chiquita, Argentina. Aquatic Conservation: Marine and Freshwater Ecosystems, 16(4), 335–347.

Mariano-Jelicich, F., Botto, F., Martinetto, P., Iribarne, O., Favero, M. (2008). Trophic segregation between sexes in the Black Skimmer revealed through the analysis of stable isotopes. Marine Biology, 155, 443–450.

Martinetto, P., Iribarne, O., Palomo, G. (2005). Effect of fish predation on intertidal benthic fauna is modified by crab bioturbation. Journal of Experimental Marine Biology and Ecology, 318, 71–84.

Martínez, S., Del Río, C. (2005). Las ingresiones marinas del Neógeno en el sur de Entre Ríos (Argentina) y Litoral Oeste de Uruguay y su contenido malacológico. Miscelánea, 14, 13–26.

Meldahl, K.H. (1993). Geographic gradients in the formation of shell concentrations: Plio-pleistocene marine deposits, Gulf of California. Palaeogeography, Palaeoclimatology, Palaeoecology, 101(1-2), 0–25.

Pereira, L. G., Fornari, M., Erthal, F., Leme, J. M., Giannini, P. C. F. (2021). Multivariate taphonomic analysis of mollusk shell concentrations in Holocene deposits of southern Brazil: An integrated approach. Palaeogeography, Palaeoclimatology, Palaeoecology, 562, 110085.

Pisano, M. F., Halpern, K. (2009). La historia de la Tierra contada desde el sur del Mundo. Geología Argentina. Ministerio de Educación. Presidencia de la Nación. Ciudad Autónoma de Buenos Aires.

Saleuddin, A. S. M., Petit, H. P. (1983). The mode of formation and the structure of the periostracum. In The mollusca (pp. 199–234). Academic Press.

Schneider-Storz, B., Nebelsick, J. H., Wehrmann, A., Federolf, C. M. J. (2008). Comparative taphonomy of three bivalve species from a mass shell accumulation in the intertidal regime of North Sea tidal flats. Facies, 54, 461–478.

Seitz, R., Lipcius, R., Olmstead, N., Seebo, M., Lambert, D. (2006). Influence of shallow water habitats and shoreline development on abundance, biomass, and diversity of benthic prey and predators in Chesapeake Bay. Marine Ecology Progress Series, 326, 11–27.

Short, A. D. (1996). The role of wave height, period, slope, tide range and embaymentisation in beach classifications: a review. Revista chilena de historia natural, 69(4), 589–604.

Silvestri, S., D’Alpaos, A., Nordio, G., Carniello, L. (2018). Anthropogenic Modifications Can Significantly Influence the Local Mean Sea Level and Affect the Survival of Salt Marshes in Shallow Tidal Systems. Journal of Geophysical Research: Earth Surface, 123, 996–1012.

Travers, A. (2007). Low-energy beach morphology with respect to physical setting: a case study from Cockburn Sound, Southwestern Australia. Journal of Coastal Research, 23(2), 429–444.

Tsolakos, K., Katselis, G., Theodorou, J. A. (2021). Taphonomy of mass mollusc shell accumulation at Amvrakikos Gulf lagoon complex sandy barriers (NW Greece). Oceanología, 63(2), 179–193.

Vila-Concejo, A., Gallop, S. L., Largier, J. L. (2020). Sandy beaches in estuaries and bays. In Sandy beach morphodynamics (pp. 343–362). Elsevier.

Violante, R. A., Cavallotto, J. L. (2004). Evolution of the semi-enclosed basins and surrounding coastal plains adjacent to the Pampean region, Argentina. Polish Geological Institute Special Papers, 11, 59–70.

Westoby, M. J., Brasington, J., Glasser, N. F., Hambrey, M. J., Reynolds, J. M. (2012). ‘Structure-from-Motion’ photogrammetry: A low-cost, effective tool for geoscience applications. Geomorphology, 179, 300–314.

Zar, J. H. (1999). Biostatistical Analysis. 4th edition. Prentice-Hall, Inc., Englewood Cliffs, NJ, 718 pp.

Zuschin, M., Stanton, R. J. (2001). Experimental Measurement of Shell Strength and its Taphonomic Interpretation. Palaios, 16(2), 161–170.

